# Early perturbations to fluid homeostasis alter development of hypothalamic feeding circuits with context-specific changes in ingestive behavior

**DOI:** 10.1101/2024.10.25.620307

**Authors:** Serena R. Sweet, Jessica E. Biddinger, Jessie B. Zimmermann, Gina L. Yu, Richard B. Simerly

## Abstract

Drinking and feeding are tightly coordinated homeostatic events and the paraventricular nucleus of the hypothalamus (PVH) represents a possible node of neural integration for signals related to energy and fluid homeostasis. We used *TRAP2;Ai14* and Fos labeling to visualize neurons in the PVH and median preoptic nucleus (MEPO) responding to both water deprivation and hunger. Moreover, we determined that structural and functional development of dehydration-sensitive inputs to the PVH from the MEPO precedes those of agouti-related peptide (AgRP) neurons, which convey hunger signals and are known to be developmentally programmed by nutrition. We also determined that osmotic hyperstimulation of neonatal mice led to enhanced AgRP inputs to the PVH in adulthood, as well as disruptions to ingestive behaviors during high-fat diet feeding and dehydration-anorexia. Thus, development of feeding circuits is impacted not only by nutritional signals, but also by early perturbations to fluid homeostasis with context-specific consequences for coordination of ingestive behavior.

## BACKGROUND/INTRODUCTION

An essential role of the hypothalamus is to coordinate multiple physiological functions and ensure homeostasis in the face of dynamic environmental changes and behavioral requirements. Eating and drinking represent tightly coordinated activities controlled by the hypothalamus^1–6^. Water restriction leads to a robust reduction in food intake, which is rapidly reversed when water is presented^7–10^. In addition to dehydration-anorexia, prandial thirst illustrates the close coordination between feeding and drinking; during food intake drinking is initiated to regulate fluids lost during meal consumption^11–13^. Such coordination requires effective integration of sensory signals conveyed to the hypothalamus by neurons that respond directly to a variety of physiological signals related to metabolic state or fluid homeostasis^14–16^.

How this integration is accomplished is not fully known but likely involves convergence of neuroendocrine sensory pathways onto common hypothalamic targets. The paraventricular nucleus of the hypothalamus (PVH) represents such a target region and plays a central role in regulating both fluid and energy balance^1,3,6,17,18^. The PVH receives direct inputs from neurons containing orexigenic agouti related peptide (AgRP) in the arcuate nucleus of the hypothalamus (ARH) that convey negative valence signals associated with hunger^19–24^. PVH neurons also receive signals related to thirst or osmotic dehydration from the median preoptic nucleus (MEPO) located in the lamina terminalis^14,25,26^. Information about hyperosmotic state is conveyed through the blood via activation of neurons in the organosum vasculosum of the lamina terminalis (OVLT) and subfornical organ (SFO), and this information is relayed to the MEPO to drive water intake^27–32^. Accordingly, Fos labeling is elevated in the PVH of osmotically dehydrated animals and in response to optogenetic activation of MEPO neurons^31,33,34^. Hunger signals resulting from fasting, or injections of 2-deoxyglucose that mimic energy deficit, also induce Fos labeling in similar components of the PVH^35,36^. Moreover, calcium imaging studies identified molecularly defined ensembles of PVH neurons that respond to both drinking and food intake^6^. Together these observations suggest that the PVH is a likely site for convergence of signals related to thirst and hunger^30,37,38^.

Development of convergent hypothalamic pathways is largely unknown, however, projections from the ARH are formed during the first two weeks of postnatal life, a time of considerable developmental plasticity^19,39–42^. AgRP neurons extend their axons from the ARH to innervate the PVH primarily during the second week of life, and postnatal perturbations in metabolic stimuli, such as leptin secretion, disturb targeting of AgRP axons in the PVH^43–49^. Moreover, manipulating the activity of AgRP neurons impairs the neurotrophic action of leptin on AgRP projections to the PVH, and offspring of dams fed a high fat diet just during lactation have impaired innervation of the PVH by AgRP and are predisposed to obesity in adulthood^50–52^. Such environmental effects on development of feeding circuitry is considered developmental programming, but how hypothalamic circuits conveying thirst signals develop has received little attention^40,41,53–56^. Neonatal mice rely on milk from dams for both nutrition and hydration, but it remains unclear if they are motivated by thirst, hunger or stimuli associated with maternal attachment^57–60^. Injections of hypertonic saline induce Fos labeling in the SFO, OVLT, and PVH of 2 day old rats, indicating that pathways for osmotic dehydration signals are intact at an early age^34^. In contrast, leptin injections do not activate PVH neurons until after they have received significant innervation from the ARH and begin transitioning to consumption of solid food^19,58,61^.

In order to identify neurons that receive convergent thirst and hunger signals in the PVH, we used adult TRAP2/Ai14 mice to permanently label PVH neurons that respond to thirst combined with Fos labeling to visualize neurons that respond to fasting^62^. In addition, we mapped the ontogeny of neuronal activation in the MEPO and PVH resulting from acute hypernatremia to visualize cells responding to osmotic dehydration. We determined that projections from the MEPO to the PVH precede those arriving from the ARH and that daily exposure to hyperosmotic stimuli during early postnatal life leads to enhanced innervation of the MEPO and PVH by AgRP neurons in adult male mice, which is associated with context-specific changes to drinking and feeding.

## RESULTS

### Drinking and feeding signals converge in the MEPO and PVH of adult mice

Water deprivation induces Fos labeling in substantial populations of neurons located in both the MEPO and PVH, and food deprivation induces robust Fos expression in the ARH and PVH^33,8^. To simultaneously visualize neurons activated by both feeding and drinking, we used the FosTRAP transgenic mouse line, *TRAP2*, in which *2A-iCreER^T2^* was knocked into the *Fos* locus to create an in-frame fusion. In *TRAP2;Ai14* double transgenic mice, neuronal activation results in the expression of CreER, which enters the nucleus in response to a 4-hydroxytamoxifen (4-OHT) injection and causes recombination, resulting in permanent expression of tdTomato in active neurons^63^. Injecting 4-OHT into *TRAP2;Ai14* double transgenic mice resulted in a marked induction of tdTomato+ labeling in the MEPO (Figure 1A-1C) and PVH (Figure 1D-1G) of mice deprived of water for 36 hours (Thirst-TRAP) compared to euhydrated controls (EUH-TRAP).

**Figure 1.**
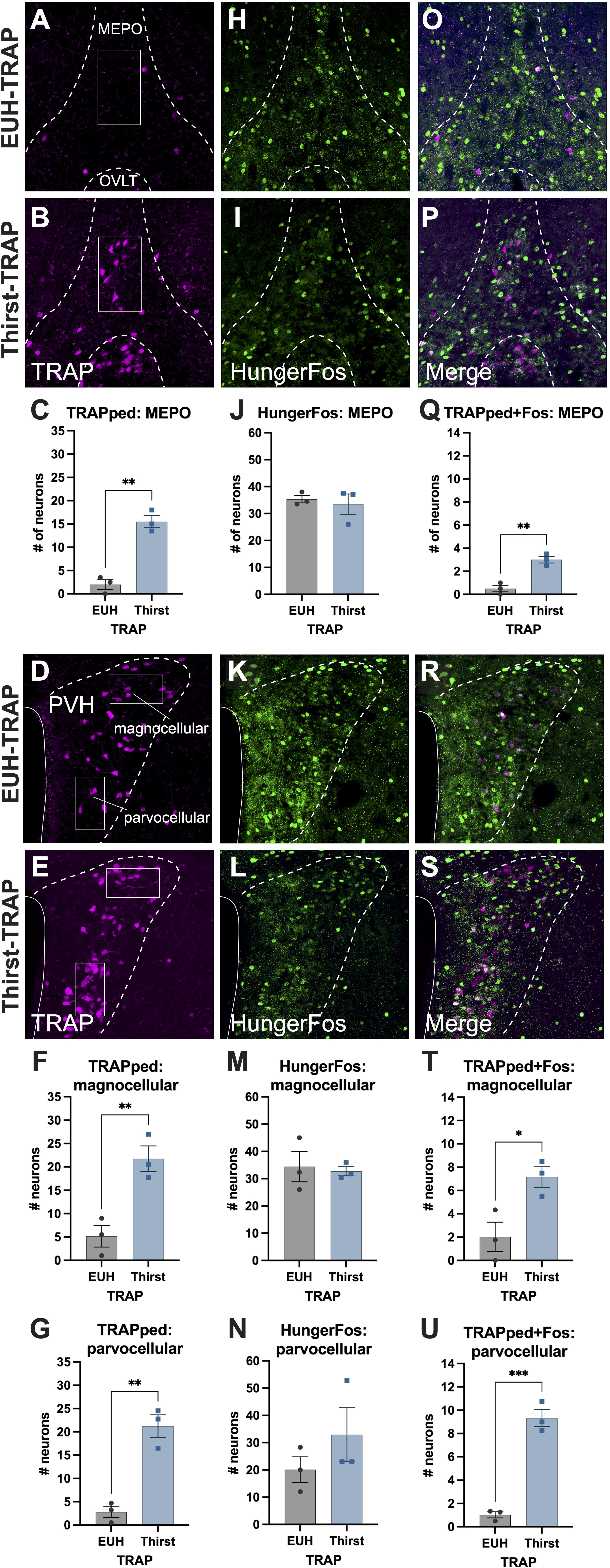
Drinking and feeding signals converge in the MEPO and PVH of adult mice. (A-B) Representative confocal images showing TRAPped (tdTomato+) cells in the MEPO of both EUH-TRAP (A) and Thirst-TRAP (B) mice. A region of interest (ROI) is indicated by the white rectangle. (C) Quantification of number of cells expressing tdTomato+ in the MEPO ROI. (D-E) Representative confocal images showing TRAPped (tdTomato+) cells in the PVH of both EUH-TRAP (D) and Thirst-TRAP (E) mice. ROIs within the magnocellular and parvocellular compartments are indicated by the white rectangles. (F) Quantification of cells expressing tdTomato+ in the magnocellular ROI of the PVH. (G) Quantification of cells expressing tdTomato+ in the parvocellular ROI of the PVH. (H-I) Representative confocal images showing Fos+ cells in the MEPO in response to a fast-refeed stimulus (Hunger-Fos) in both EUH-TRAP (H) and Thirst-TRAP (I) mice. (J) Quantification of immunohistochemical (IHC) analysis of cells expressing Fos+ in the MEPO ROI. (K-L) Representative confocal images showing Fos+ cells in the PVH in response to a fast-refeed stimulus (Hunger-Fos) in both EUH-TRAP (K) and Thirst-TRAP (L) mice. (M) Quantification of IHC analysis of cells expressing Fos+ in the magnocellular ROI of the PVH. (N) Quantification of IHC analysis of cells expressing Fos+ in the parvocellular ROI of the PVH. (O-P) Representative confocal images showing co-labeled tdTomato+/Fos+ cells in the MEPO in both EUH-TRAP (O) and Thirst-TRAP (P) mice. (Q) Quantification of IHC analysis of co-labeled tdTomato+/Fos+ cells in the MEPO. (R-S) Representative confocal images showing co-labeled tdTomato+/Fos+ cells in the PVH in both EUH-TRAP (R) and Thirst-TRAP (S) mice. (T) Quantification of IHC analysis co-labeled tdTomato+/Fos+ cells in the magnocellular compartment of the PVH. (U) Quantification of IHC analysis of co-labeled tdTomato+/Fos+ cells in the parvocellular compartment of the PVH. All data are represented as mean ± SEM and data points are quantified across 1-2 sections for individual animals. Thirst-TRAP (n = 3), and EUH-TRAP (n = 3). Unpaired t-test; * p < 0.05, **p <0.01, *** p < 0.001. Abbreviations: MEPO, median preoptic nucleus; OVLT, vascular organ of the lamina terminalis; PVH, paraventricular nucleus of the hypothalamus. See also Figure S1.

Thirst-TRAPped cells in the PVH were not domain specific, with the magnocellular and parvocellular cells of the PVH exhibiting a significantly higher induction in Thirst-TRAP mice compared to EUH-TRAP controls. As expected, there were no differences in tdTomato+ labeling in the ARH in Thirst-TRAP mice compared to EUH-TRAP controls (S1A-S1C). After Thirst-TRAP, we challenged all mice with a 23 hour fast followed by ad-libitum refeeding 1 hour before perfusion and tissue collection (Hunger-Fos). There were no differences in Hunger-Fos+ neurons in the MEPO (Figure 1H-1J), PVH (Figure 1K-1N), and ARH (Figure S1D-S1F) of EUH-TRAP and Thirst-TRAP groups following fast-refeeding. When looking at co-labeled cells, there was significantly more Hunger-Fos co-labeled Thirst-TRAPped cells in the MEPO compared to EUH-TRAP controls (Figure 1O-1Q). Additionally, there was significantly greater Hunger-Fos co-labeling in Thirst-TRAPped cells in both the magnocellular and parvocellular compartments of the PVH (Figure 1R-1U). We did not detect double-labeled neurons in ARH (Figure S1G-S1I). Together, these results suggest dehydration signals in adult mice activate sub-populations of neurons in the MEPO and PVH, feeding signals activate sub-populations of neurons in the MEPO, PVH, and ARH, and convergence of these signals is detected in the MEPO and PVH.

### MEPO neurons project to the PVH during the first week of life

Although direct projections from the MEPO to the PVH are known to be important for fluid homeostasis in adults, whether these projections are in place in neonatal mice has yet to be established. To visualize the MEPO projections to the PVH in neonatal mice, we used DiI axonal labeling in wild-type neonatal mice perfused at P8 (Figure 2A). Single DiI crystals were implanted in the MEPO (Figure 1B-1D) using an insect needle and brains were incubated in fixative for 6 weeks. We observed a moderate density of labeled axons extending caudally from the MEPO and remaining largely confined to the periventricular zone of the hypothalamus. Moreover, MEPO neurons appear to provide at least a moderate density of DiI labeled axons in the PVH (Figure 2E-2G). These results suggest that neural projections from the MEPO to the PVH are intact in early postnatal life.

**Figure 2.**
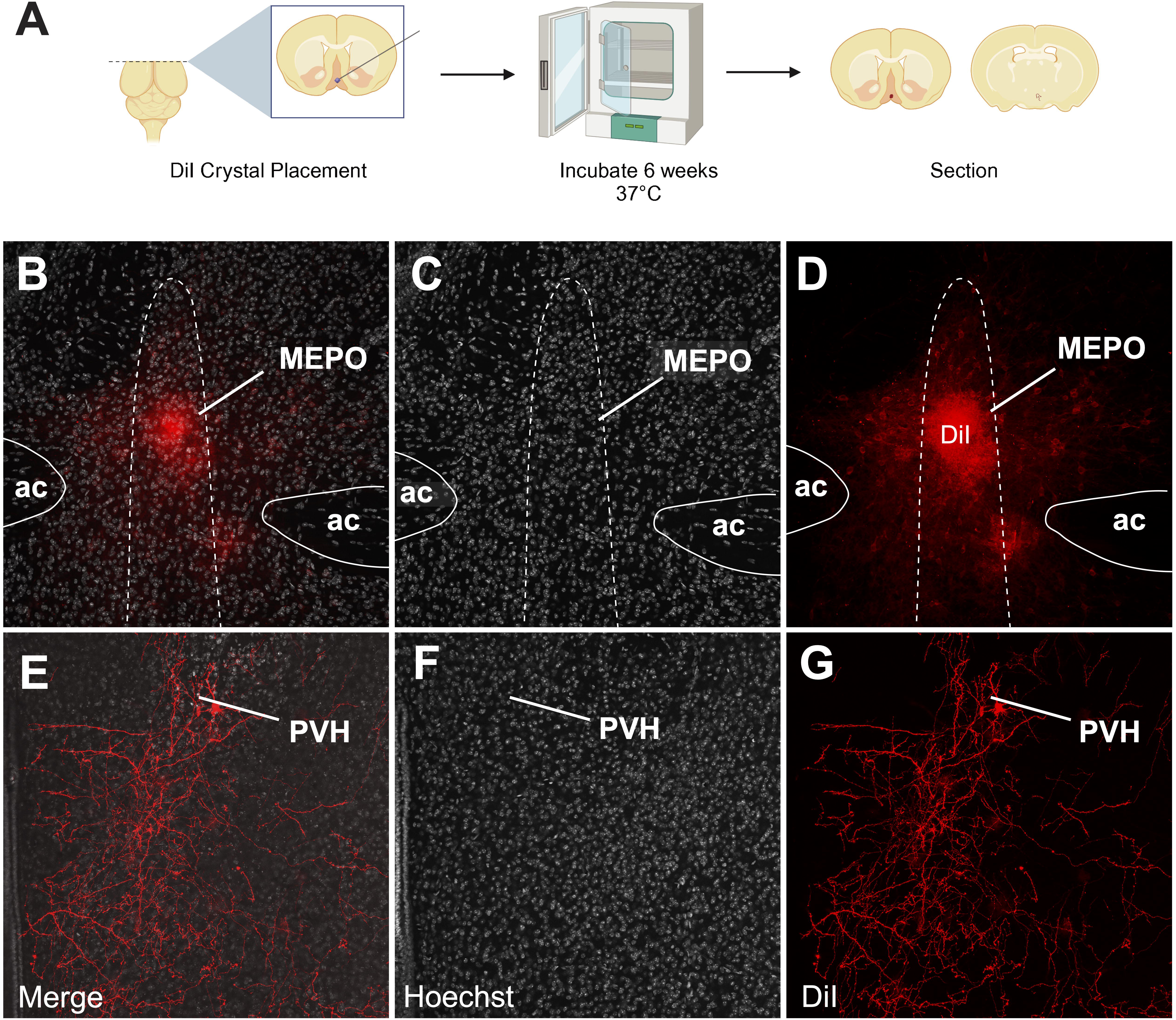
Postnatal developmental projections from the MEPO to the PVH. (A) Schematic of experiment design, showing brains from P8 male mice were sectioned from rostral to caudal so as to expose the MEPO and single DiI crystals were implanted in the MEPO using an insect needle, the brains were then placed back into fixative and incubated for 5-6 weeks, and then sectioned on a vibratome. (B-D) Confocal images showing Hoechst Dye and implant location of DiI crystal in the MEPO. (E-G) Confocal images showing Hoechst Dye and downstream robust DiI axonal labeling in the PVH.

### Ontogeny of MEPO and PVH neurons responding to osmotic stimuli

The DiI labeling results establish that neurons in the MEPO project to the PVH early in postnatal life, but whether these neurons are capable of responding to dehydration signals and conveying this information to the PVH early in postnatal life in mice is unknown. Therefore, we investigated development of the functional capacity of projections from the MEPO to the PVH by visualizing Fos immunolabeling in response to a dehydration stimulus. Subcutaneous (s.c.) administration of 0.1 mL/10g BW of 2.0M NaCl (hypertonic saline, HS), or of 0.9% NaCl (isotonic saline, IS), were given to male mice at P1, P8, P16, and P30. Significant increases in Fos labeling were observed in the MEPO at P1, P8, P16, and P30 in HS-treated mice compared to IS controls (Figure 3A-3L), indicating that MEPO neurons are capable of responding to osmotic signals at birth. Moreover, significantly more HS-induced Fos-immunoreactive nuclei were detected in the PVH at P1, compared to controls, suggesting that dehydration signals are conveyed from the MEPO to the PVH as early as the first postnatal day (Figure 3M, 3Q, 3U). There is also significant induction of Fos in the PVH at P8, P16, and P30 compared to controls (Figure 3N-3P, 3R-3T, 3V-3X). Notably, the maximum density of Fos labeling in the MEPO was observed by P8 (Figure 3Y). This contrasted with the later achievement of maximal levels of Fos labeling in the PVH at P16 (Figure 3Z), which coincides with maturation of projections from AgRP neurons. Together, these results suggest MEPO and PVH neurons are responding to osmotic signals as early as P1, with a graded maximal response coinciding with the early divergence of feeding and drinking behavior.

**Figure 3.**
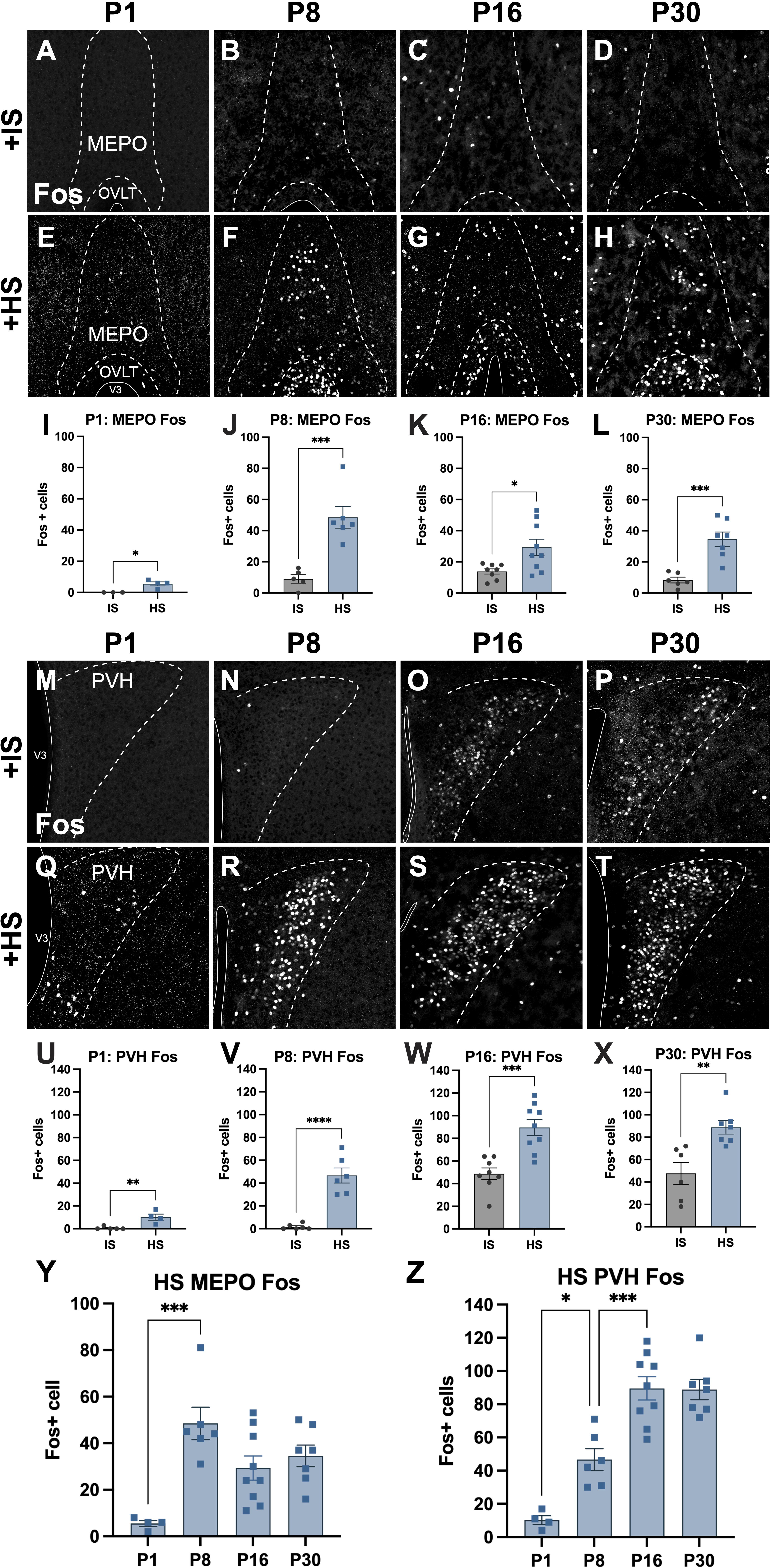
Ontogeny of circuits responding to osmotic stimuli. (A-D) Representative confocal images showing Fos+ cells in the MEPO of neonatal mice across ages P1 (A), P8 (B), P16 (C), and P30 (D) treated with a subcutaneous (s.c.) administration of 0.1mL/10g BW of 0.9% NaCl (isotonic saline, IS) as a control. (E-H) Representative confocal images showing Fos+ cells in the MEPO of neonatal mice at ages P1 (E), P8 (F), P16 (G), and P30 (H) treated with s.c. administration of 0.1mL/10g BW of 2.0M NaCl as a dehydration stimulus (hypertonic saline, HS). (I) Quantification of IHC analysis of Fos+ cells in the MEPO at P1. IS (n = 3), and HS (n = 4). (J) Quantification of IHC analysis of Fos+ cells in the MEPO at P8. IS (n = 5), and HS (n = 6). (K) Quantification of IHC analysis of Fos+ cells in the MEPO at P16.IS (n = 8), and HS (n = 9). (L) Quantification of IHC analysis of Fos+ cells in the MEPO at P30. IS (n = 6), and HS (n = 7). (M-P) Representative confocal images showing Fos+ cells in the PVH of neonatal mice across ages P1 (M), P8 (N), P16 (O), and P30 (P) treated with a s.c. administration of 0.1mL/10g BW of 0.9% NaCl as a control (IS). (Q-T) Representative confocal images showing Fos+ cells in the PVH of neonatal mice at ages P1 (Q), P8 (R), P16 (S), and P30 (T) treated with s.c. administration of 0.1mL/10g BW of 2.0M NaCl as a dehydration stimulus (HS). (U) Quantification of IHC analysis of Fos+ cells in the PVH at P1. IS (n = 5) and HS (n = 4). (V) Quantification of IHC analysis of Fos+ cells in the PVH at P8. IS (n = 6) and HS (n = 6). (W) Quantification of IHC analysis of Fos+ cells in the PVH at P16. IS (n = 8) and HS (n = 9). (X) Quantification of IHC analysis of Fos+ cells in the PVH at P30. IS (n = 6) and HS (n = 7). (Y) Comparison of Fos+ cells in the MEPO of HS mice across ages. One-way ANOVA with multiple comparisons. (Z) Comparison of Fos+ cells in the MEPO of HS mice across ages. One-way ANOVA with multiple comparisons. All data are represented as mean ± SEM and data points are quantified across 1 section for individual animals. * p < 0.05, **p <0.01, *** p < 0.001.

### Postnatal perturbation of fluid homeostasis impacts development of AgRP projections to the MEPO and PVH

To determine if activation of neurons during osmotic dehydration alters innervation of the PVH by AgRP neurons, we adapted an HS-injection protocol used previously in adult mice to model developmental disruption in fluid homeostasis by administering postnatal (PN) daily injections from P5-P15 of hypertonic saline (HS^PN^) or isotonic saline (IS^PN^) and then assessed development of AgRP inputs to the MEPO and PVH in adulthood^29^. Compared to IS^PN^ control mice, the density of AgRP axons in HS^PN^ mice is significantly elevated in the MEPO (Figure 4A-4C) and in the PVH (Figure 4D-4F). Notably, treatment of adult HS^PN^ mice with acute injections of HS (150ul of 2.0M NaCl) before tissue collection did not result in altered levels of Fos induction in the MEPO (Figure 4G-4I) or PVH (Figure 4J-4L) compared to that observed in IS^PN^ mice, which suggests that the postnatal HS treatments do not have a sustained adverse effect on osmotic signaling from the MEPO to the PVH. Together, these results suggest that development of AgRP projections is impacted by the stimulation of neural circuits involved in drinking during postnatal life, without deleterious effects on the function of drinking circuitry in adulthood.

**Figure 4.**
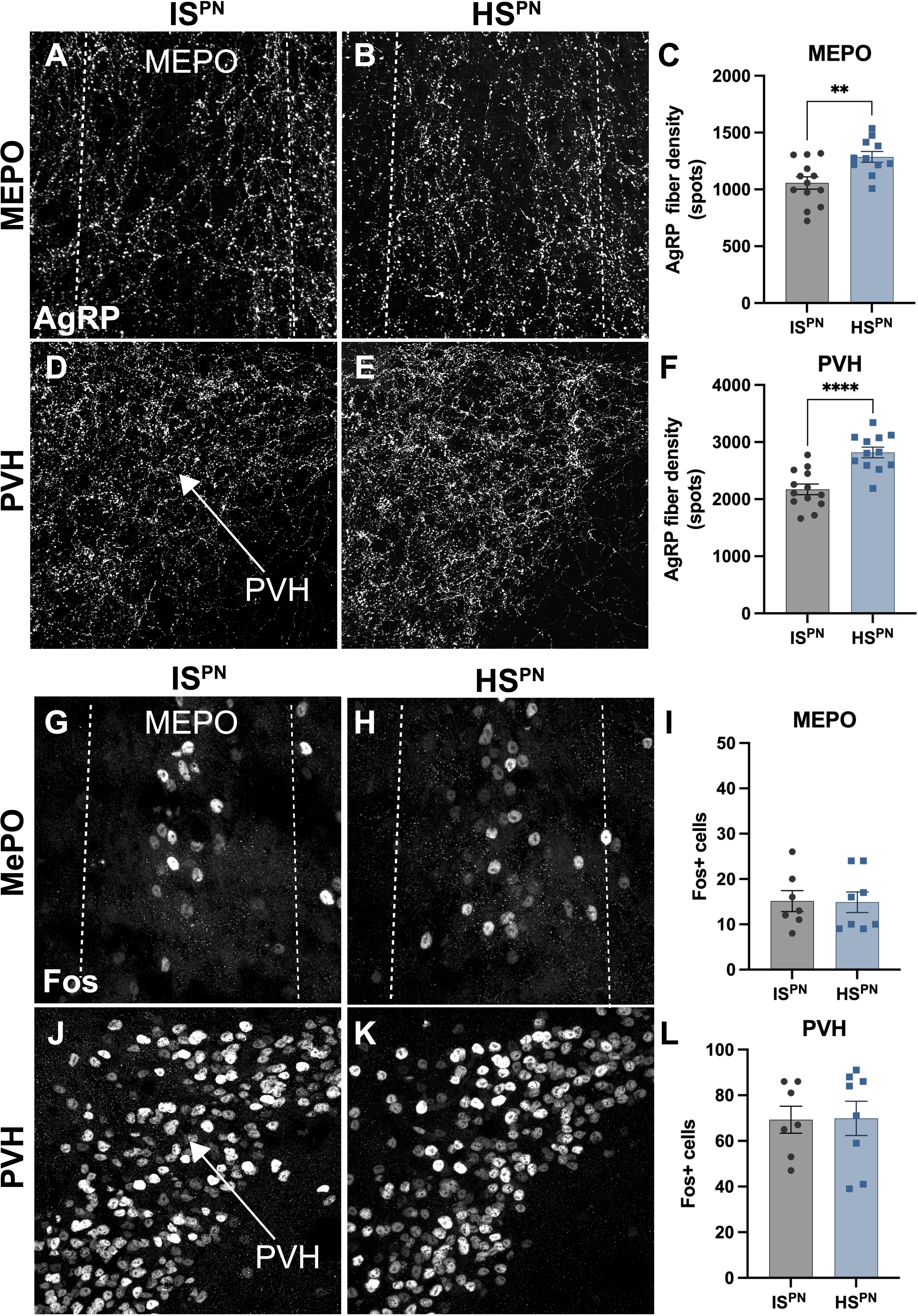
Postnatal perturbation to fluid homeostasis impacts development of AgRP projections to the MEPO and PVH. (A-B) Representative confocal images showing AgRP fiber densities in the MEPO of adult mice treated daily from P5-P15 with either s.c administration of 0.9% NaCl as a control (IS^PN^; A) or 2.0M NaCl as a dehydration stimulus (HS^PN^, B). (C) Quantification of IHC analysis of AgRP fiber densities in the MEPO using the spots feature in Imaris software. IS^PN^ (n = 13) and HS^PN^ (n = 11). (D-E) Representative confocal images showing AgRP fiber densities in the PVH of IS^PN^ (D) and HS^PN^ (E) mice. (F) Quantification of IHC analysis of AgRP fiber densities in the PVH using the spots feature in Imaris software. IS^PN^ (n = 13), and HS^PN^ (n = 12). (G-H) Representative confocal images showing Fos+ cells in the MEPO of IS^PN^ (G) and HS^PN^ (H) mice administered an acute intraperitoneal (i.p.) injection of 250ul 2.0M NaCl (HS) 90 minutes before tissue collection. (I) Quantification of IHC analysis of Fos+ cells in the MEPO administered HS i.p. 90 minutes before tissue collection. IS^PN^ (n = 7), and HS^PN^ (n = 8). (J-K) Representative confocal images showing Fos+ cells in the PVH of IS^PN^ (J) and HS^PN^ (K) mice administered HS i.p. injection 90 minutes before tissue collection. (L) Quantification of IHC analysis of Fos+ cells in the PVH administered HS i.p. 90 minutes before tissue collection. IS^PN^ (n = 7), and HS^PN^ (n = 8). All data are represented as mean ± SEM and data points are quantified across 1 section for individual animals. Unpaired t-test; * p < 0.05, **p <0.01, *** p < 0.001.

### Hyperstimulation of drinking circuits during postnatal development leads to increases in water intake during HFD feeding in adults

Because osmotic dehydration altered the density of AgRP inputs to the PVH, we examined ingestive behavior in mice exposed to HS during the perinatal period. A cohort of adult HS^PN^ and IS^PN^ mice were singly housed in metabolic profiling chambers with ad libitum access to standard chow (normal chow diet, NCD) and water for 5 days, followed by data collection for 7 days with access to high-fat diet (HFD). No differences in body weight were observed between groups (S2A) and HS^PN^ mice did not display a difference in water or food intake compared to IS^PN^ controls fed NCD (Figure S2B-S2C). Moreover, HS^PN^ mice did not display a significant change in HFD food intake compared to IS^PN^ controls (Figure 5A-5B). However, HS^PN^ mice displayed a significant increase in cumulative water intake during HFD exposure (Figure 5C-5D). These data suggest that early perturbations to fluid homeostasis have a lasting effect on ingestive behavior that appears to be diet-dependent.

**Figure 5.**
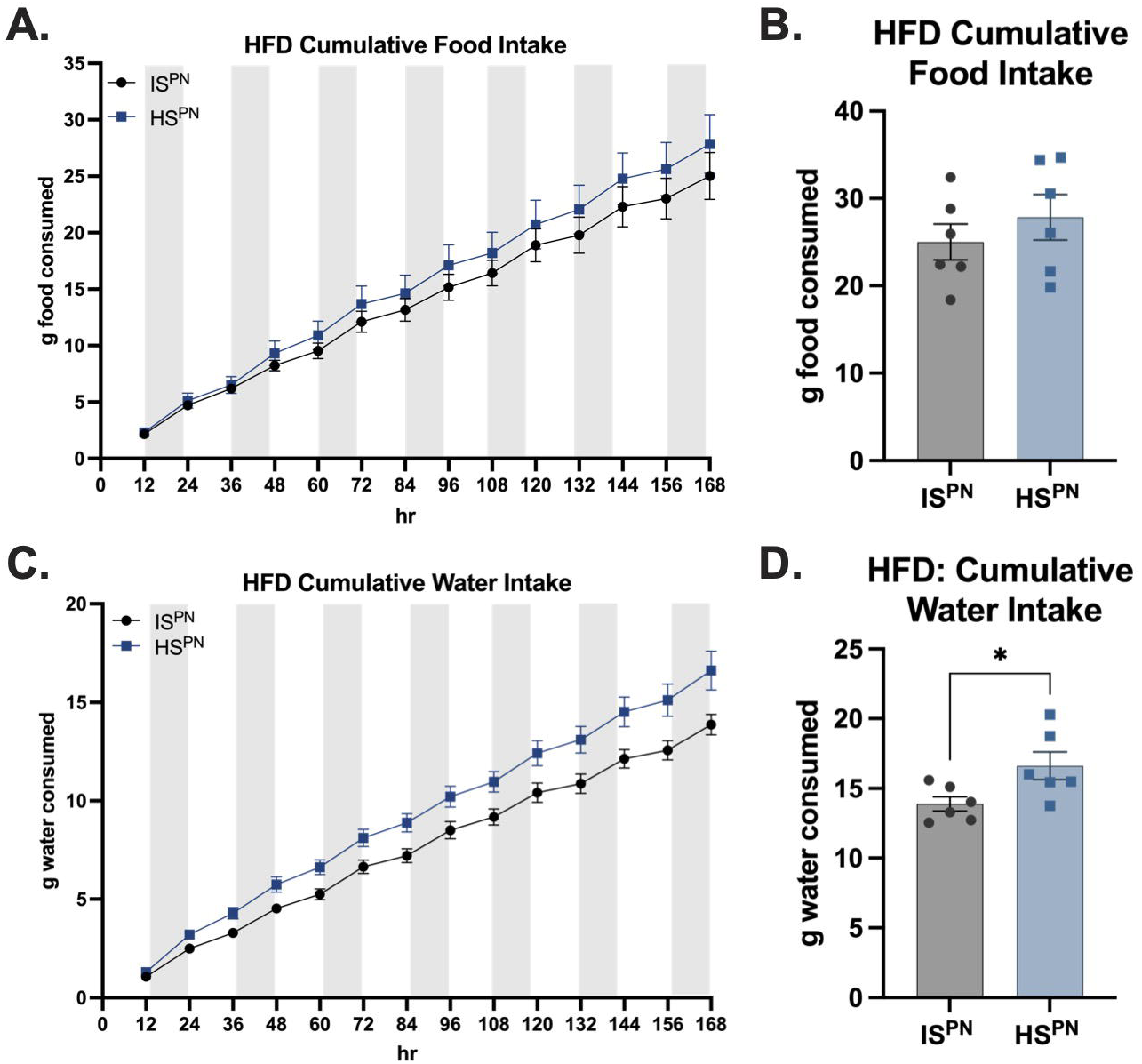
Hyperstimulation of drinking circuits during postnatal development increases water intake in the context of a HFD. (A) Quantification of cumulative food intake during the 168hrs (7 days) of HFD exposure in IS^PN^ (n = 6) and HS^PN^ (n = 6) animals. 2-way ANOVA: no significant effect of treatment over time (p = 0.7423). Data are presented as group mean values ± SEM. (B) Comparison of final cumulative food intake during HFD exposure. IS^PN^ (n = 6) and HS^PN^ (n = 6). (C) Quantification of cumulative water intake during the HFD exposure. IS^PN^ (n = 6) and HS^PN^ (n = 6). 2-way ANOVA: a significant main effect of treatment and time (p < 0.0001). Data are presented as group mean values ± SEM. (D) Comparison of final cumulative water intake during HFD exposure. IS^PN^ (n = 6) and HS^PN^ (n = 6). Data in (B, D) are represented as mean ± SEM and data points are individual animals. Unpaired t-test; *p < 0.05, **p <0.01, *** p < 0.001. See also Figure S2.

### Hyperstimulation of drinking circuits during postnatal development leads to a sustained dehydration-anorexic response in adults

Because feeding and drinking are linked behaviors, we adapted a model of dehydration-anorexia to determine if HS^PN^ mice show normal anorexic responses to dehydration in adulthood^7,9,10,36^. A cohort of adult HS^PN^ and IS^PN^ mice were singly housed in metabolic profiling chambers with ad libitum access to food and water for a 4-days, followed by a 48-hr dehydration period, and a subsequent 4-day period of rehydration. During the light phase, HS^PN^ mice did not display a change in food or water intake during the acclimation, dehydration, or rehydration periods (Figure 6A, Figure S3A). Although HS^PN^ mice did not display a significant change in food intake before or during dehydration (Figure 6B-6D), there was a significant and sustained decrease in food intake during the first dark cycle after rehydration (Figure 6E). There were no significant changes in water intake before or after the dehydration period in HS^PN^ mice (Figure S3B-S3D). Together, these results suggest that osmotic dehydration during postnatal development permanently perturbs patterns of ingestion with context-specific changes, revealing perturbations to fluid homeostasis as a novel effector of developmental programming.

**Figure 6.**
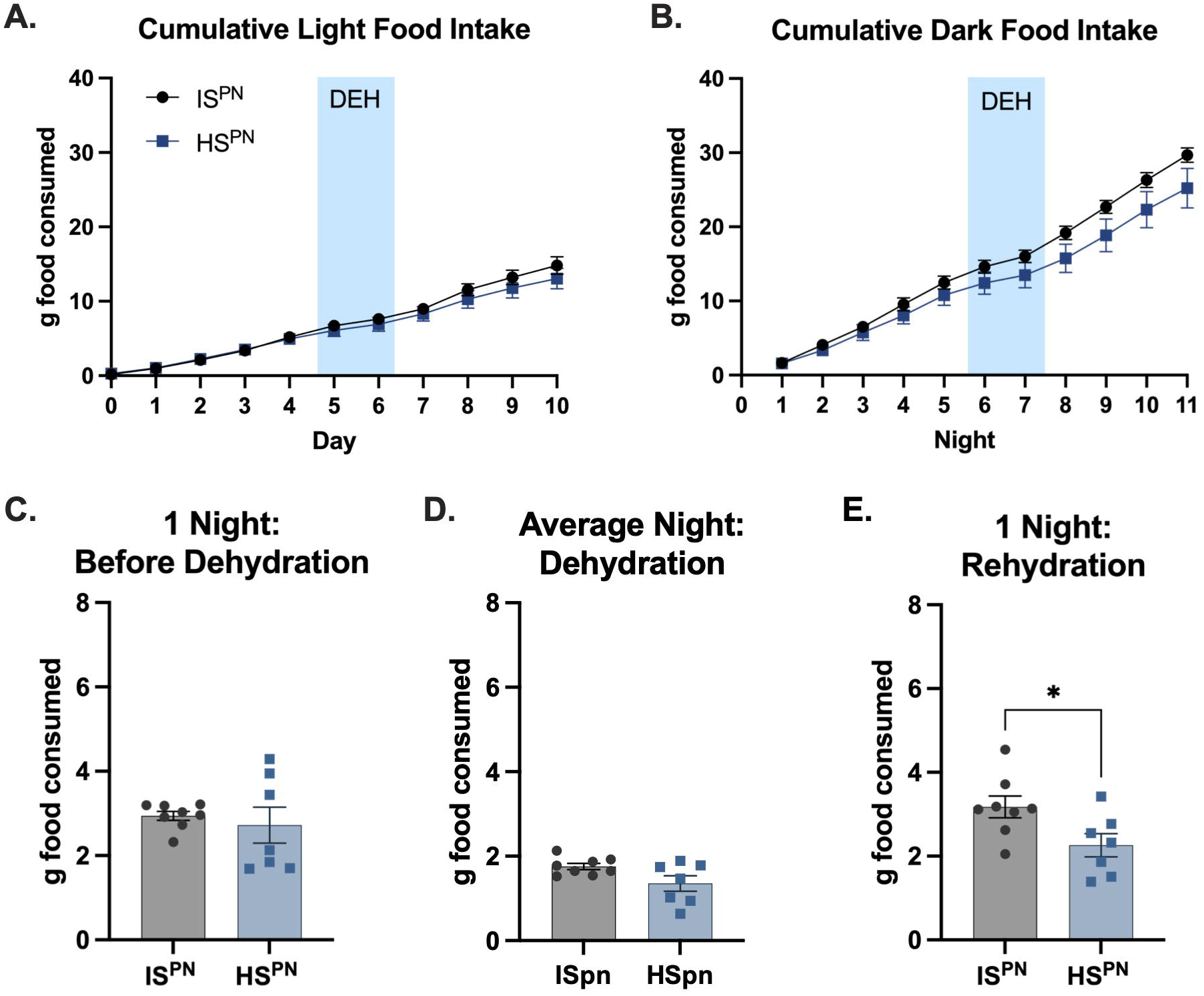
Hyperstimulation of drinking circuits during postnatal development leads to a sustained dehydration-anorexic response in adult mice. (A) Quantification of cumulative food intake during the inactive light cycle of IS^PN^ mice (n = 8) and HS^PN^ mice (n = 7). 2-way ANOVA: no significant effect of treatment over time (p = 0.3310). Data are presented as group mean values ± SEM. (B) Quantification of cumulative food intake during the active dark cycle of IS^PN^ (n = 8) and HS^PN^ (n = 7). 2-way ANOVA: a significant main effect of treatment over time (p = 0.0028). Data are presented as group mean values ± SEM. (C) Quantification of food intake the night before the dehydration period. IS^PN^ (n = 8) and HS^PN^ (n = 7). (D) Quantification of average food intake for the two nights of the dehydration period. IS^PN^ (n = 8) and HS^PN^ (n = 7). (E) Quantification of food intake for the first night following return of water to the cage. IS^PN^ (n = 8) and HS^PN^ (n = 7). Data for (C-F) are represented as mean ± SEM and data points are individual animals. Unpaired t-test; *p < 0.05, **p <0.01, *** p < 0.001. See also Figure S3.

## DISCUSSION

Neural connections between hypothalamic nuclei develop primarily during postnatal life and appear to be sensitive to alterations in a variety of environmental factors during this period^43–49^. Our results indicate that projections from MEPO neurons to the PVH are established early in postnatal life and precede development of AgRP inputs to the PVH from the ARH. As early as the first day of life (P1) signals conveying osmotic stimuli activate neurons in the MEPO and PVH. However, peak levels of Fos labeling in the PVH appear to be delayed, suggesting that HS-induced neuronal activation in the PVH requires complete innervation by the MEPO to fully respond to such an osmotic stimulus. We also observed enhanced densities of AgRP terminals in the PVH of mice that received hyperstimulation of thirst circuits by postnatal osmotic dehydration. This observation suggests that afferents to the PVH from osmotically activated MEPO neurons may influence development of convergent AgRP afferents, and that such circuitry defects could correspond to observed context-specific changes in ingestive behavior. Together, these results suggest a new model of developmental programming that links postnatal exposure to hyperosmotic dehydration stimuli and development of feeding circuits.

### Drinking and feeding signals converge in the MEPO and PVH of adult mice

Our thirst and hunger double-labeling results in adult TRAP2 mice demonstrate that a subpopulation of neurons located in a restricted domain of the PVH represents a cellular mode of action for convergence of signals conveying thirst signals with those related to hunger. While this is the first clear demonstration that individual neurons respond to both thirst and feeding signals, previous evidence from both rat and mouse models identified the PVH as a likely node of sensory integration for coordination of drinking and food intake^36^. Although it is well established that osmotic dehydration signals are sensed by the SFO and OVLT and then conveyed to the MEPO and PVH, ablation of MEPO or PVH, but not the circumventricular organs, abolishes dehydration-induced drinking^17,31^. Similarly, signals from AgRP neurons convey hunger signals to the PVH that are required for normal regulation of feeding, but the presence of Fos labeling in the MEPO in fasted mice suggests that a subpopulation of these neurons may also respond to both thirst and hunger ^18,22,64^. The number of neurons in the PVH we detected that are double-labeled likely represent a conservative estimate for numbers of neurons capable of responding to convergent sensory stimuli. Because Fos labeling identifies only those neurons with detectable Fos expression at the time of perfusion, it is unlikely that the density of labeling detected represents maximum numbers of neurons activated by hunger. In contrast, the tdTomato labeling resulting from activity-dependent expression of Cre-recombinase identifies neurons receiving the cumulative effects of dehydration. Nevertheless, the high density of double-labeled tdTomato+/Fos+ cells in the MEPO and PVH responding to sequential dehydration and fasting indicate that feeding and drinking signals are linked at a cellular level and supports a model in which ensembles of PVH and MEPO neurons encode a combinatorial representation of cues related to both fluid and energy homeostasis to coordinate drinking and feeding behavior.

### Ontogeny of neurons in the MEPO and PVH activated by osmotic signals aligns with development of structural connectivity

Using postnatal treatment of HS to model dehydration at different timepoints, we demonstrate functional connectivity during early postnatal development where the MEPO and PVH display Fos labeling in response to dehydration signals as early as P1, consistent with earlier studies in the rat where there is robust Fos induction in the MEPO in response to HS at P2^34^. Furthermore, the ontogeny of functional connectivity is evidenced by the progressive increases in Fos labeling in response to HS, where the maximum Fos induction we observe in the MEPO and PVH is at the end of the first and second week of life, respectively. This finding is consistent with the structural connectivity between the MEPO and PVH at P8 visualized with DiI labeling. Similarly, AgRP neurons do not appear to drive milk consumption until after the second week of life, coinciding with the exploration of solid food and the maturation of projections from AgRP neurons to their targets, including the PVH^19,57,60^. Thus, development of projections from the MEPO to the PVH corresponds to the ability of osmotic signals to activate the PVH neurons and precedes the functional maturation of AgRP inputs from the ARH.

### Postnatal perturbation of fluid homeostasis impacts the development of AgRP projections to the MEPO and PVH

Developing hypothalamic neural circuits are vulnerable to environmental perturbations during postnatal critical periods^39,40,43^. Leptin-deficiency, suppressed AgRP activity, and maternal high fat diet during the postnatal critical period for development of AgRP neurons all result in reduced innervation of downstream neuronal targets^19,47,48,50^. In contrast, daily exposure to HS leads to an increase in AgRP innervation of the PVH and MEPO, suggesting that hyperstimulation of MEPO neurons promotes innervation of common targets by AgRP neurons. Whether the hyperinnervation of PVH neurons by AgRP afferents is due to a transsynaptic activation of PVH neurons or is due to activation of local guidance cues that promote innervation by AgRP neurons remains to be determined. However, the greater sensitivity of MEPO neurons to HS stimulation during early postnatal life is consistent with the possibility that observed changes in structural connectivity could be due to activity-dependent development mechanisms, similar to those observed previously for AgRP inputs to the PVH, and associated leptin-dependent development of projections from the PVH oxytocin neurons to the brainstem^52^. Together these findings suggest a model in which projections from AgRP neurons to the PVH develop during postnatal life under the influence of both cell autonomous activity, as well as that of activation of convergent inputs to PVH neurons derived from MEPO neurons. For example, stimulation of glutamatergic afferents to the PVH by MEPO neurons during development may lead to enhanced activity of PVH neurons that promotes convergent innervation by GABAergic AgRP neurons. Alternatively, stimulation of the MEPO may promote development of a descending glutamatergic input to the ARH, where hyperstimulation of AgRP neurons during development may lead to increased convergent innervation of PVH neurons. Although it will take additional experimentation to distinguish between these possible developmental mechanisms, our data demonstrate that dehydration signals acting on the MEPO alter development of feeding circuitry and provide a novel example of developmental programming in the context of early dehydration.

### Hyperstimulation of drinking circuits during postnatal development leads to context-specific changes in ingestive behaviors

In addition to observed changes in AgRP projections caused by postnatal exposure to osmotic dehydration, we also observed sustained changes to ingestive behavior displayed by HS^PN^ offspring in adulthood. During HFD exposure, adult HS^PN^ mice showed enhanced water intake with no change in food intake. Moreover, dehydration of adult HS^PN^ mice led to a sustained anorexic response after water replacement with no changes to water intake, in sharp contrast to the normal reversal of dehydration-anorexia that follows water replacement in control mice. Thus, postnatal osmotic dehydration appears to disrupt the tight coupling of fluid homeostasis and food intake and appears to be context-specific. These findings also suggest that HS^PN^ may impact neuronal integration of signals controlling fluid homeostasis and hunger related to competing motivational states. One possible explanation is that changes to convergent circuits conveying thirst and hunger signals may alter the impact of negative valence signals that represent environmental cues associated with nutrient or water ingestion^24,29,65^. This interpretation is consistent with the observation that drinking was only perturbed in HS^PN^ mice during exposure to HFD and that food intake was only altered within the context of dehydration-anorexia. Alternatively, observed changes in ingestive behavior could be due to defects in anticipatory modulation, consistent with other findings that water intake changes with diet content^11,13^. Thus, metabolic information being conveyed to the MEPO may not be appropriately integrated in HS^PN^ mice due to structural changes in the neural circuits involved in feeding, resulting in changes to ingestive behavior during metabolic challenge. Collectively, our results support a model in which dehydration in early life not only leads to long-term structural changes in hypothalamic circuits, but also expression of context-specific functional deficits in coordinated ingestive behaviors^66^.

## Conclusion

Drinking and feeding are highly coordinated behaviors, and we have identified a novel developmental interaction between neurons that sense osmotic challenges and AgRP circuits that may contribute to environmentally-specified programming of context-specific ingestive behavior. The developmental activity demonstrated here for osmotic dehydration during postnatal life not only changes patterns of structural connectivity in the hypothalamus but exerts a lasting change to how fluid balance and food intake are coordinated during homeostatic stress. Thus, convergent neural inputs to the PVH from neurons in the MEPO and ARH may represent a developmental substrate that links early dehydration with sustained perturbation of fluid homeostasis and energy balance, adding osmotic disruptors to the growing list of agents that act during critical periods of development to program hypothalamic structural and functional connectivity. As water scarcity and dietary hypernatremia increase in vulnerable populations, this developmental exposure may have unforeseen consequences for how the brain coordinates multiple aspects of homeostasis.

## LIMITATIONS OF THE STUDY

While dehydration appears to be a novel model of developmental programming, additional work is required to define molecular mechanisms mediating the effects of postnatal dehydration on development of AgRP innervation patterns. It will also be informative to define specific cell types visualize with double-labeled Thirst-TRAP/Hunger-Fos cells in the PVH. Although our purpose here was to use development of projections from AgRP neurons to represent a programmable component of feeding circuits, they are only one of many cell types involved in feeding. For example, evaluating changes in POMC neurons to the PVH or afferents to PVH neurons that express melanocortin receptors will provide additional insight into changes imposed onto feeding circuitry by osmotic thirst during postnatal life. The PVH clearly serves as a node of integration for signals important for coordination of ingestive behavior, but there may be other components of feeding circuitry that are equally responsive to developmental programming by postnatal dehydration. For example, the lateral hypothalamus (LHA) has been implicated as a site of integration of feeding and drinking, highlighting the importance of defining the organization and development of assessing a broader set of regions involved in processing information regulating fluid and energy homeostasis. Nevertheless, our studies provide a foundation for future discovery into how multiple environmental signals can exert a lasting effect on the functional anatomy of ingestive homeostasis.

## ACKNOWLEDGEMENTS

This research was supported by the Eunice Kennedy Shriver National Institute of Child Health and Human Development (5F31HD106890) and the National Institute of Diabetes and Digestive and Kidney Diseases (5R01DK106476).

## AUTHOR CONTRIBUTIONS

S.R.S. and R.B.S. conceived the project, designed the experiments, and wrote the manuscript. S.R.S. performed Fos induction, AgRP immunolabeling, and ingestive behavior experiments. S.R.S. performed statistical analysis and preparation of all figures. S.R.S and J.E.B. performed TRAP experiments with Fos immunolabeling. J.E.B assisted in figure preparation of TRAP data. J.B.Z and G.L.Y. assisted in Fos and AgRP immunohistochemical experiments and data analysis for neonatal and adult mice. R.B.S. led funding acquisition and supervision.

## STAR METHODS

### Key resources table

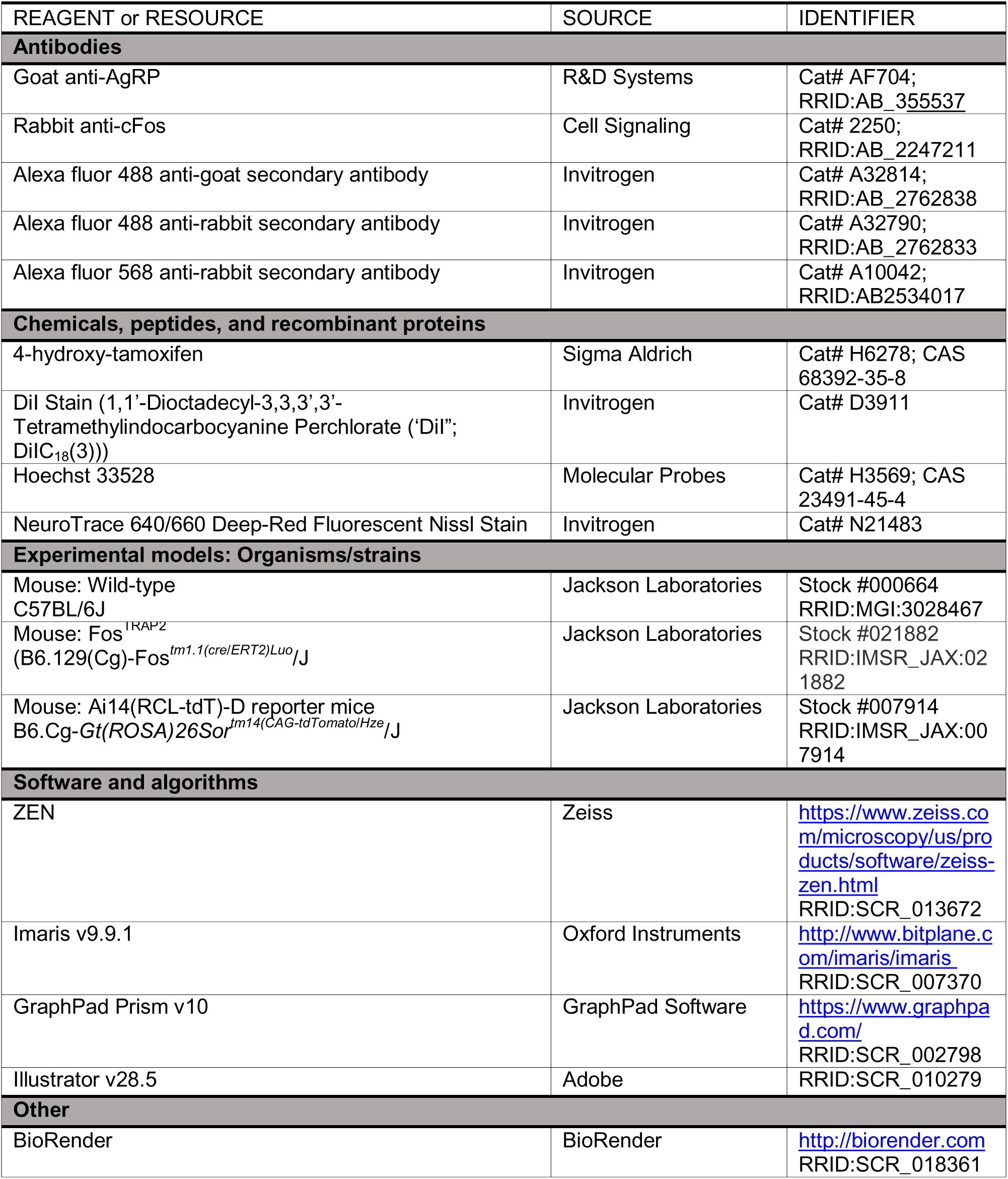

## RESOURCE AVAILABILITY

### Lead contact

Further information and requests for resources and reagents should be directed to and will be fulfilled by the lead contact, Richard B. Simerly

### Materials availability

This study did not generate new unique reagents.

### Data and code availability

- All data supporting the findings of this study will be shared by the lead contact upon request.
- This paper does not report original code.
- Any additional information required to reanalyze the data reported in this paper is available from the lead contact upon request.

## EXPERIMENTAL MODEL AND SUBJECT DETAILS

### Animals

All animal care and experimental procedures were performed in accordance with the National Institutes of Health Guide for the Care and Use of Laboratory Animals (Garber et. al, 2011) and approved by the Institutional Care and Use Committee at Vanderbilt University. Mice were housed at 22°C on a 12 hr light-dark cycle and provided ad libitum access to water and standard chow diet (PicoLab Rodent Diet 20, #5053), unless otherwise noted. Mice were weaned at P21 and maintained with mixed genotype littermates until males were used for experiments.

Wild-type (C57BL/6J; stock #000664) male mice at P1, P8, P16, and P30 were used for the acute dehydration and DiI experiments and at P48-60 for postnatal dehydration experiments. Male adult (8-12 weeks of age) Fos^TRAP2^ (B6.129(Cg)-Fos*^tm1.1(cre/ERT2)Luo^*/J; stock #021882) mice expressing an inducible Cre recombinase were crossed with Ai14(RCL-tdT)-D reporter mice (B6.Cg-*Gt(ROSA)26Sor^tm14(CAG-tdTomato/Hze^*/J; stock #007914) to induce Cre-dependent expression of the tdTomato fluorescent protein in neurons activated by treatment with hypertonic saline. Fos^TRAP2^ mice were heterozygous for both the *Fos^2A-iCreER^* and Ai14 alleles. All mice were obtained from the Jackson Laboratory.

## METHOD DETAILS

### Tissue Preparation

Mice were perfused and processed for histochemical experiments as previously described (Biddinger et al., 2020; Bouyer & Simerly 2013), unless otherwise specified. Briefly, mice were first anesthetized with tribromoethanol (TBE), then perfused transcardially with 0.9% saline followed by ice-cold fixative (4% paraformaldehyde in 0.1M borate buffer, pH 9.5) for 20 min. Brains were carefully removed from the skull and postfixed in the same fixative for 4 hr at 4°C.

For immunohistochemical studies, brains were cryoprotected overnight in 20% sucrose and a sliding microtome was used to collect 30μm thick coronal sections from brains derived from adult mice (P48-P60), or 50μm thick sections from brains collected at postnatal ages (P1-P30). The free-floating sections were stored in cryoprotectant solution at -20°C until further processing.

For DiI tracing studies, brains were stored in fixative at 4°C until DiI implantation. To prepare the brains for DiI implantation, the brains were mounted in a 3% agarose solution made in 0.02M KPBS. Brains were mounted onto a vibratome stage and sectioned from rostral to caudal so as to expose the median preoptic nucleus (MEPO). Single DiI crystals were implanted into the MEPO using an insect needle, and the brains were placed back into fixative and incubated for 5-6 weeks at 37°C. After incubation, brains were sectioned on a vibratome at 80μm.

### Immunohistochemistry

To prepare the tissue for immunohistochemical processing, tissue sections were removed from cryoprotectant and rinsed in 0.02M KPBS. Sections were incubated overnight at 4°C in a 0.02M KPBS blocking buffer containing 2% normal donkey serum (Jackson ImmunoLabs) and 0.3% Triton X-100. Primary antibodies were diluted in blocking buffer, and used at the following concentrations: goat anti-AgRP (1:1000; R&D Systems) and rabbit anti-cFos (1:1000; Cell Signaling). Tissue sections were incubated in blocking buffer containing combinations of primary antibodies for 48 hr at 4°C. Sections were rinsed in 0.02M KPBS, and then incubated in the appropriate species-specific fluorophore-conjugated Alexa Fluor secondary antibodies for 1 hr at RT. Tissue sections were rinsed again in 0.02M KPBS and counterstained with Hoechst Dye and/or NeuroTrace, and rinsed in 0.02M KPBS. Tissue sections were mounted directly onto charged Superfrost Plus slides (Fisher Scientific), and after drying overnight at 4°C were coverslipped with No.1.5 Gold Seal cover glass (Electron Microscopy Services) and ProLong Gold antifade mounting medium (Invitrogen).

### DiI tracing studies

Free-floating tissue sections from the vibratome were counterstained at 1:10000 with Hoechst Dye in 0.02M KPBS. Tissue sections were rinsed in 0.02M KPBS and then mounted directly onto charged Superfrost Plus slides and coverslipped with glycerol buffer.

### cFos experiments

cFOS immunostaining was used to assess location of activated neurons in response to dehydration and food deprivation. Briefly, age-matched groups of mice received subcutaneous (s.c.) injections of sterile hypertonic saline (2.0M NaCl) or isotonic saline (0.9% NaCl) on P1, P8, P16, or P28 for the postnatal studies; intraperitoneal (i.p.) injections of 250μL of 2.0M NaCl (P48-P80) for IS/HS^PN^ anatomical studies; and a 23 hr fast followed by a refeed was used for the Hunger-Fos paradigm. Mice were anesthetized with TBE 60 min later, perfused, and tissue processed for immunohistochemistry as described above.

### Postnatal hypertonic saline treatments

Wild-type mouse litters were adjusted to 6-8 pups. Neonatal mice were weighed daily and remained in their home cage with their littermates and biological dam, with ad libitum access to standard rodent chow and water. To activate neuronal circuits involved in drinking via osmotic challenge, mice received daily subcutaneous (s.c.) injections from P5-P15 of either hypertonic saline (2.0M NaCl, HS^PN^) or isotonic saline (0.9% NaCl, IS^PN^). Injections were administered in the home cage, at least 1 hr prior to lights out. Mice were then weaned onto the same chow diet at P21 and maintained on chow diet into adulthood. Upon completion of physiological testing in adulthood, mice were perfused and processed for immunohistochemistry as described above.

### TRAP induction

Adult *TRAP2;Ai14* mice were singly housed the week prior to utilization, and experiments were carried out in the homecage. All mice underwent a handling acclimation procedure daily, with an i.p. injection of 0.9% NaCl, for five days prior to the Thirst-TRAP experiment. 4-hydroxytamoxifen (4-OHT) was dissolved at 20 mg/mL in ethanol by shaking at 37°C for 15 min and the dissolved 4-OHT was then stored at -20°C until used. Before use, the dissolved 4-OHT was mixed with corn oil, shaken at 37°C, and ethanol evaporated for a final concentration of 10 mg/mL. For Thirst-TRAP studies, the final 10 mg/mL 4-OHT solution was injected i.p. at a dose of 50 mg/kg (Allen et. Al 2017) 30 hr after water bottles were removed from the cage. Water was returned 6 hr after 4-OHT injection. Mice were carefully observed for signs of distress or ill health during the water deprivation period. To identify neurons activated by hunger (Hunger-Fos), after 1 week of recovery from Thirst-TRAP, food was removed from the homecage. Mice were fasted for 23 hr before food was replaced. 60 min after refeeding, mice were perfused and processed for immunohistochemical localization of Fos induction, as described above.

### Food and water intake analysis

All mice used in Promethion Core System (Sable Systems International) studies, unless otherwise stated, were adults treated daily from P5-P15 with either hypertonic saline (HS^PN^; 2.0M NaCl) or isotonic saline (IS^PN^; 0.9% NaCl) as described above. As adults, mice were placed in individual cages the week prior to Sable testing, with ad libitum access to standard rodent chow and water. Mice were then singly housed in Promethion cages placed inside a temperature-controlled cabinet for the duration of the metabolic experiments described below and returned to their respective homecages following the conclusion of each experiment.

For the high-fat diet challenge, mice were singly housed in Promethion cages and acclimated to the temperature-controlled cabinet for collection of baseline metabolic measurements for 5 days. A high-fat diet (40%, Research Diets) was then introduced for 7 days with continuous measurement of food and water intake.

For the dehydration-anorexia challenge, mice were singly housed in Promethion cages and acclimated to the temperature-controlled cabinet for collection of baseline food and water intake for 4 days. The water bottle was then removed for 48 hr, and the mice were carefully observed for signs of distress or ill health every 8 hr during the water deprivation period. The water bottle was then re-introduced into the cage and continuous measurements of food and water intake were collected for 4 days.

## QUANTIFICATION AND STATISTICAL ANALYSIS

### Image acquisition and analysis

Brain tissue sections containing the MEPO, PVH, and ARH were identified and examined on a laser scanning confocal microscope (Zeiss 800). Cytoarchitectonic features of these nuclei and subnuclei were visualized and identified with nuclear Hoechst Dye and/or cytoplasmic NeuroTrace to define matching regions of interest (ROI) for quantitative analysis. Anatomically defined ROIs were used to quantify the density of labeled AgRP axons as well as cFOS in the MEPO, PVH, and ARH. In order to visualize and quantify AgRP axon densities to the PVH and MEPO of adult mice, sections were imaged at high magnification using an oil-immersion 40x objective, at a frequency of 0.08μm in the x and y planes and z-step of 0.427. All other images were captured with a 20x objective and collected at a frequency of 1.18μm in the x and y planes, and a z-step of 1.38μm, with a digital zoom of 0.8x unless otherwise noted.

Imaris (Bitplane V9.0) visualization software was used to prepare 3D reconstructions of each multichannel set of images. To quantify the overall densities of labeled AgRP fibers in the ROI of the MEPO and PVH, the spots function in Imaris was used to segment the images and measure the density of AgRP terminals within the ROI. To quantify the number of Fos nuclei in the MEPO and PVH induced by HS injection or Thirst-TRAP, the number of Fos nuclei was counted in a maximum projection of confocal images through each ROI of each region, guided by the spots function in Imaris software.

### Statistical analysis and graphical display of data

Statistical analyses were performed using GraphPad Prism software (Version 9.5). Food and water intake data were analyzed using repeated-measures ANOVA tests to compare data between groups over multiple time points; unpaired t-tests were used to compare data for all other statistical tests unless otherwise noted. Data are presented as group mean values ± SEM, as well as individual data points. Differences between groups were considered statistically significant at *p*<0.05. The Median Absolute Deviation (MAD) statistical method was used to detect outliers in datasets used for food and water intake analysis, with a 3.0 threshold (Leys et. al 2013). The median is a measure of central tendency that is insensitive to the presence of outliers compared to using the mean as a measure of central tendency. Graphs were constructed with GraphPad Prism (v10; GraphPad Software). Figure layouts were organized using Illustrator (v28.5; Adobe Systems). Photoshop was also used to adjust brightness and contrast on confocal images (v25.11; Adobe Systems).

---

**Supplementary Figure 1.**
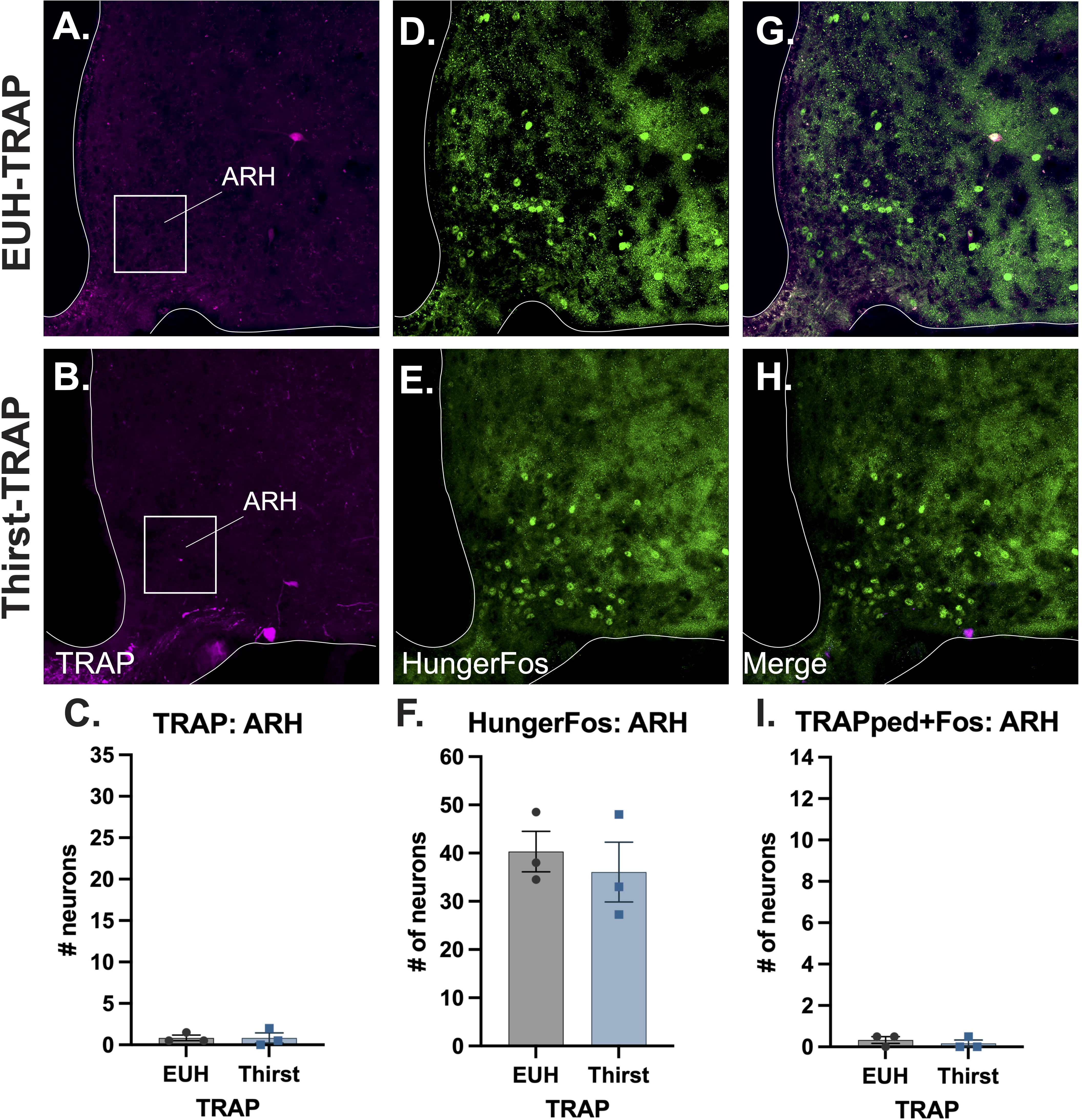
Drinking and feeding signals do not converge in the ARH of adult mice. (A-B) Representative confocal images showing TRAPped (tdTomato+) cells in the ARH of both EUH-TRAP (A) and Thirst-TRAP (B) mice. A region of interest (ROI) is indicated by the white rectangle. (C) Quantification of number of cells expressing tdTomato+ in the ARH ROI. (D-E) Representative confocal images showing Fos+ cells in the ARH in response to a fast-refeed stimulus (Hunger-Fos) in both EUH-TRAP (D) and Thirst-TRAP (E) mice. (F) Quantification of immunohistochemical (IHC) analysis Fos+ cells in the ARH ROI. (G-H) Representative confocal images showing co-labeled TdTomato+/Fos+ cells in the ARH in both EUH-TRAP (G) and Thirst-TRAP (H) mice. (I) Quantification of IHC analysis co-labeled tdTomato+/Fos+ cells in the ARH. All data are represented as mean +/- SEM and data points are quantified across 1-2 sections for individual animals. Thirst-TRAP (n = 3), and EUH-TRAP (n = 3). Unpaired t-test; * p < 0.05, **p <0.01, *** p < 0.001.

**Supplementary Figure 2.**
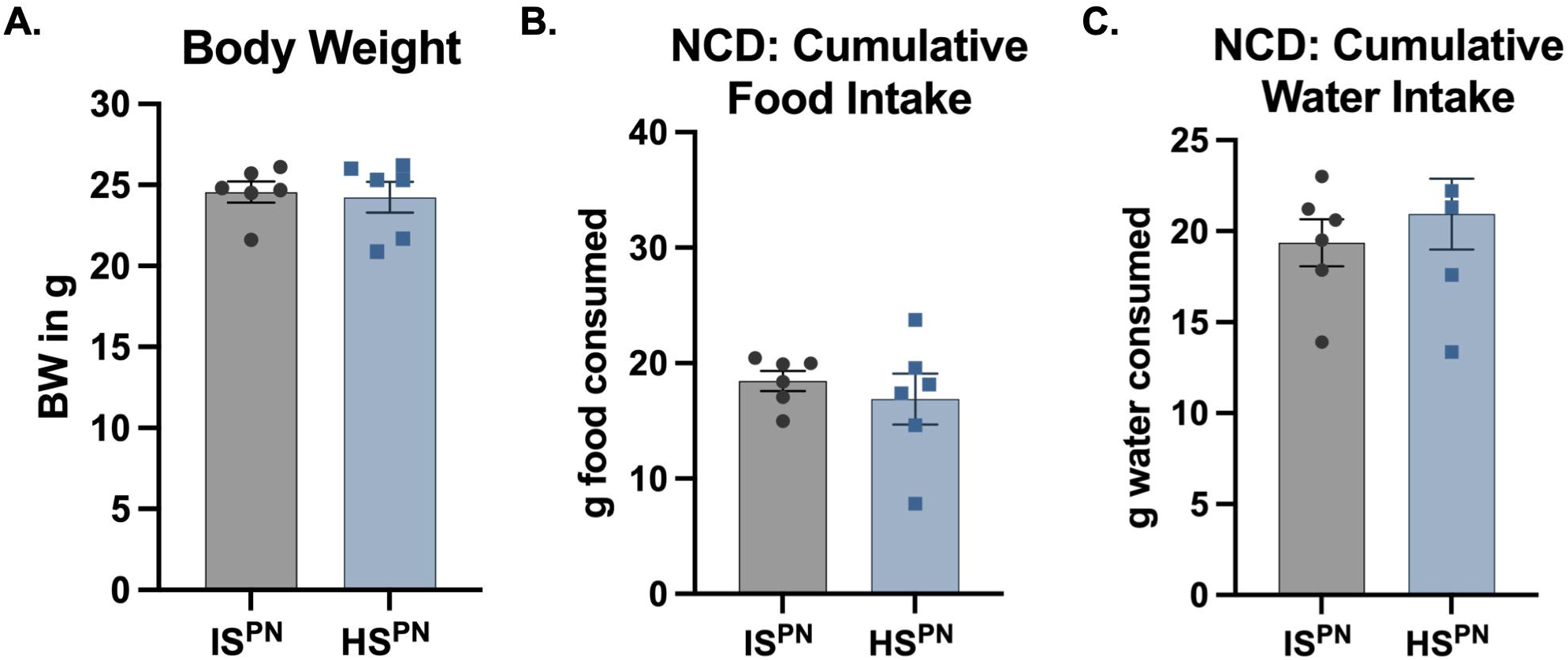
Hyperstimulation of drinking circuits during postnatal development does not affect body weight or food intake in the context of NCD. (A) Quantification of body weight of adult IS^PN^ (n = 6) and HS^PN^ (n = 6) mice. (B) Quantification of food intake during the normal chow diet (NCD). IS^PN^ (n = 6) and HS^PN^ (n = 6). (C) Quantification of water intake during the NCD. IS^PN^ (n = 6) and HS^PN^ (n = 6). All data are represented as mean +/- SEM and data points are quantified across individual animals. Unpaired t-test; * p < 0.05, **p <0.01, *** p < 0.001.

**Supplementary Figure 3.**
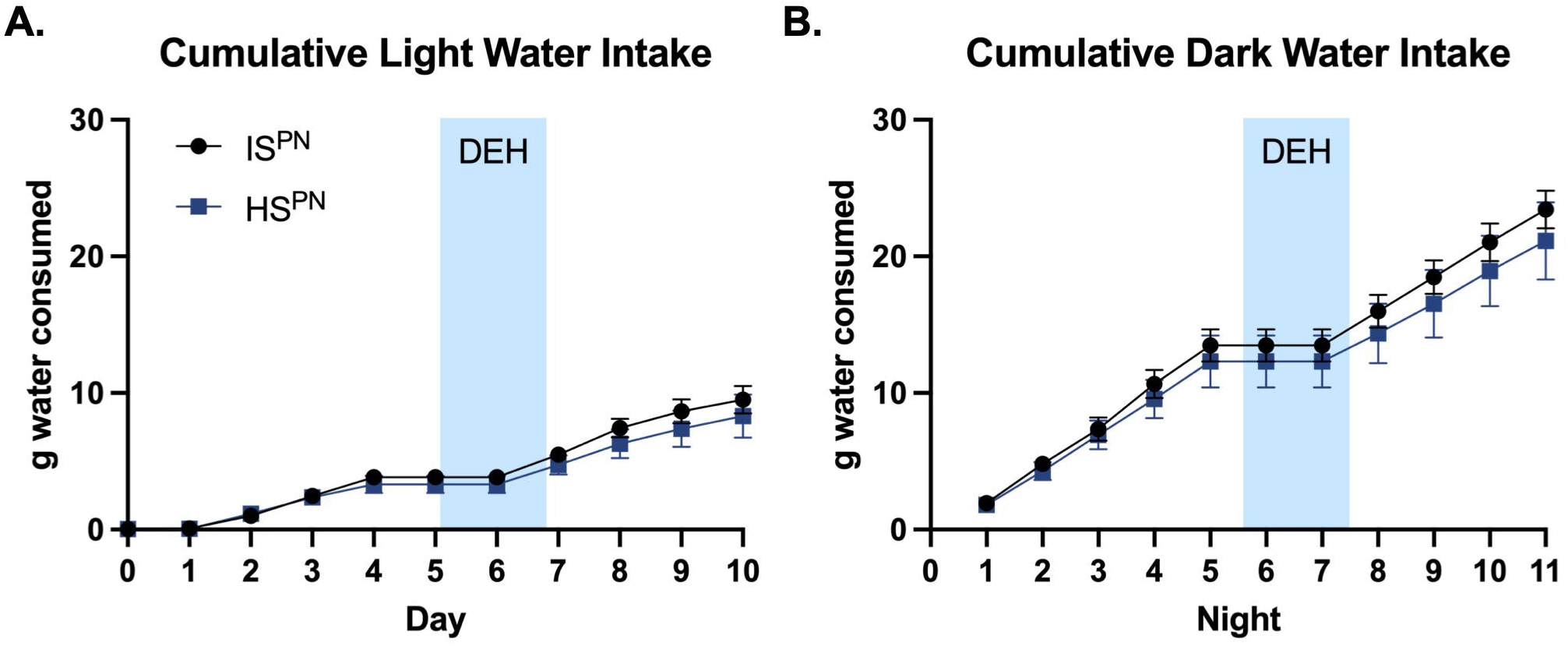
Hyperstimulation of drinking circuits during postnatal development does not change water intake before or after dehydration. (A) Quantification of cumulative water intake during the inactive light cycle in control IS^PN^ adult mice (n = 8) and HS^PN^ mice (n = 7). 2-way ANOVA: no significant effect of treatment over time (p = 0.7208). (B) Quantification of cumulative water intake during the active dark cycle. IS^PN^ (n = 8) and HS^PN^ (n = 7). 2-way ANOVA: no significant effect of treatment over time (p = 0.8952). All data are presented as group mean values ± SEM.

